# Optimisation of a multiplexed, high throughput assay to measure neutralising antibodies against SARS-CoV-2 variants

**DOI:** 10.1101/2024.09.01.610733

**Authors:** Caroline L. Ashley, Malik Bloul, Sibel Alca, Lachlan Smith, Wang Jin, David Khoury, Claudio Counoupas, Miles Davenport, James A. Triccas, Megan Steain

## Abstract

A multiplexed, lentivirus-based pseudovirus neutralisation assay (pVNT) was developed for high-throughput measurement of neutralising antibodies (nAbs) against three distinct SARS-CoV-2 spike variants. Intra-assay variability was minimised by optimising the plate layout and determining an optimal percentage transduction for the pseudovirus inoculum. Comparison of monoclonal antibody EC_50_ titres between single and multiplexed pVNT assays showed no significant differences, indicating reliability of the multiplexed assay. Evaluation of convalescent human sera confirmed assay validity, with consistent fold drops in EC_50_ for variant pseudoviruses relative to the ancestral strain observed across single and multiplexed assays. This multiplexed pVNT provides a reliable tool for assessing nAb responses against SARS-CoV-2 variants and could be used to accelerate preclinical vaccine assessment in preparation for the next coronavirus pandemic.

## Introduction

Serological assays are important tools for measuring immune responses against pathogens. The severe acute respiratory syndrome coronavirus-2 (SARS-CoV-2) pandemic drove the rapid development of a variety of assays to measure antibody responses in small animal models as well as convalescent and vaccinated individuals. SARS-CoV-2 neutralising antibody (nAb) titres strongly correlate with protection from symptomatic (Khoury et al., 2021) and severe disease (Cromer et al., 2023). However, nAbs wane over time and their ability to neutralise is decreased against emerging variants (Cromer et al., 2022). This has necessitated the ongoing development of booster vaccines. To accelerate the approval of new vaccines, immunobridging using nAb titres from pre-clinical models and/or small clinical trials has been proposed (Gilbert et al., 2022; Khoury et al., 2023; Ramasamy, 2023). However, a major hurdle to using nAbs as a correlate of protection is a lack of assay standardisation, which contributes to issues with reproducibility and variability (Khoury et al., 2023; Manak et al., 2024; Wang et al., 2020; Zheng et al., 2022). Additionally, demonstrating breadth of neutralisation against a broad array of SARS-CoV-2 variants in pre-clinical testing can be challenging given the limited blood volumes that can be obtained from small animals.

The ‘gold-standard’ for measuring nAb titres are live virus neutralisation assays, however, these are time consuming, and require a biosafety level 3 containment facility (Bain et al., 2020) as well as highly trained staff. Consequently, these assays are expensive and have limited throughput. To overcome some of these limitations, pseudotyping of replication-deficient lentiviruses or vesicular stomatitis virus with SARS-CoV-2 spike proteins has been widely adopted (Tandon et al., 2020). These pseudovirus-based systems recapitulate SARS-CoV-2 spike-mediated binding and entry and deliver a reporter gene to the cell, either luciferase or a fluorescent protein. Pseudovirus neutralisation tests (pVNTs) therefore present a safer and more cost-effective system to measure functional nAb levels. The replication-deficient nature of pseudoviruses also ensures genetic stability in the encoded spike variants being assessed, providing greater consistency across experiments compared to assays using live virus which may mutate with successive passage in cell lines (Ogando et al., 2020; Sonnleitner et al., 2022).

To efficiently screen samples against a large panel of pseudovirus variants, multiplexing of pseudovirus assays can be employed (Nie et al., 2016). This approach can also minimise variability, by running several pseudoviruses against the same antibody-containing sample on the same cells concurrently. We have developed a high-throughput multiplex pseudovirus assay that allows for assessment of nAb titres against three SARS-CoV-2 variants simultaneously from small starting volumes of sera. We examined several parameters that influence assay variability and determined optimal conditions to minimise inter-run variance. We show that nAb titres determined by our multiplexed assay correlate with those measured by single pseudovirus assays. Our multiplexed pVNT allows for rapid high-throughput screening of preclinical samples to expedite testing of new coronavirus vaccine candidates and could be easily adapted to measure neutralising antibodies against other viruses.

## Materials and Methods

### Cells and viruses

HEK293T and HEK293T.ACE2 cells were maintained in Dulbecco’s modified Eagle medium (DMEM; Life Technologies, Thermo Fisher Scientific, Waltham, MA, USA) as described previously (Norman et al., 2021). Replication-deficient pseudotyped lentivirus particles were generated by co-transfecting the vector plasmid: pBCKS(HIV-1SDmCMVeGFP-P2A-luc2pre-IIU) (Tiffen et al., 2010) or pCDH-CMV-MCS-EF1-Neo containing mTAG-BFP2 or LSS-mOrange engineered in-house, and a SARS-CoV-2 spike expression construct in pcDNA3.1 or pCAGGS backbone, with lentivirus packaging and helper constructs (Koldej et al., 2005; Tiffen et al., 2010) into HEK293T cells using Fugene HD (Promega, Madison, WI, USA). Harvested pseudovirus particles were filtered and stored at -80°C.

### Neutralisation assays

Pseudovirus used in pVNT were combined at the dilution factors previously determined by serial dilution titration to give 5% transduction (per variant). Diluted pseudovirus(es) were incubated with human serum or mAbs at various dilutions/concentrations for 1 hour at 37°C, prior to spinoculation (800 g) onto ACE2 over-expressing 293T cells. 72 hours post-transduction, cells were fixed and stained with Syto60™ Red fluorescent nucleic acid stain (Life Technologies, Thermo Fisher Scientific), as per the manufacturer’s instructions, imaged using an Opera Phenix Plus high content screening system (Revvity, Waltham, MA, USA) and the percentage of cells positive for each fluorescent protein was enumerated (Harmony® high-content analysis software, Revity). Neutralising antibody titres, the dilution or concentration required for 50% inhibition of infection (EC50), were calculated by interpolation in GraphPad Prism 10 using a 5 PL sigmoidal curve.

### Patient and murine samples

In March 2020 when ancestral SARS-CoV-2 was the dominant circulating variant in Sydney, Australia (COVIMM cohort), unvaccinated patients over age 5 with COVID-19 were recruited through the Royal Prince Alfred hospital (RPAH) COVID clinic or Virtualcare system. Blood samples were collected by an experienced staff member at 2 to 4 months after primary infection. This study protocol was approved by the RPAH ethics committee, human ethics number X20-0117 and 2020/ETH00770. Consent was provided by all participants. 25 mL of blood from each participant was collected in tubes containing lithium heparin anticoagulant (Sigma-Aldrich, Merck, Darmstadt, Germany) and centrifuged at 200 xg for 10 min at RT. The resulting top layer of plasma was isolated and centrifuged again at 1000 xg for 10 minutes. Plasma samples were heat-inactivated at 56°C for 30 min to eliminate any infectious material in accordance with the Charles Perkins Centre physical containment level-2 guidelines. Heat-inactivated plasma was then stored in 200 μL aliquots at -30°C. Murine serum samples were obtained following vaccination with an adjuvanted SARS-CoV-2 trimeric spike protein antigen.

## Results and Discussion

### Multiplexed pseudovirus neutralisation assay workflow

To develop a high throughput system to measure nAbs against multiple SARS-CoV-2 variants, we developed a multiplexed pVNT using a 384-well plate format. This assay allows for simultaneous determination of nAb titres against three distinct SARS-CoV-2 spike expressing pseudoviruses. The workflow involved is outlined in Figure 1A. To allow multiplexing, lentiviral particles pseudotyped with different SARS-CoV-2 spike variants are distinguished from one another by the fluorophore transgene they deliver, namely EGFP, mTAG-BFP2 or LSS-mOrange. When used with the nucleic acid counterstain Syto™60 these fluorophores can be detected using a high content screening system with minimal spectral interference. In wells containing the 3-pseudovirus mixture, cells co-expressing EGFP, mTAG-BFP and LSS-mOrange can be observed alongside those expressing individual fluorophores (Figure 1B). The presence of singly transduced cells suggests that mixing the pseudoviruses is unlikely to alter their entry. The 384-well plate format permits high-throughput, with low starting volumes of samples, thus maximising the data that can be obtained from small starting volumes of sera.

**Figure 1.**
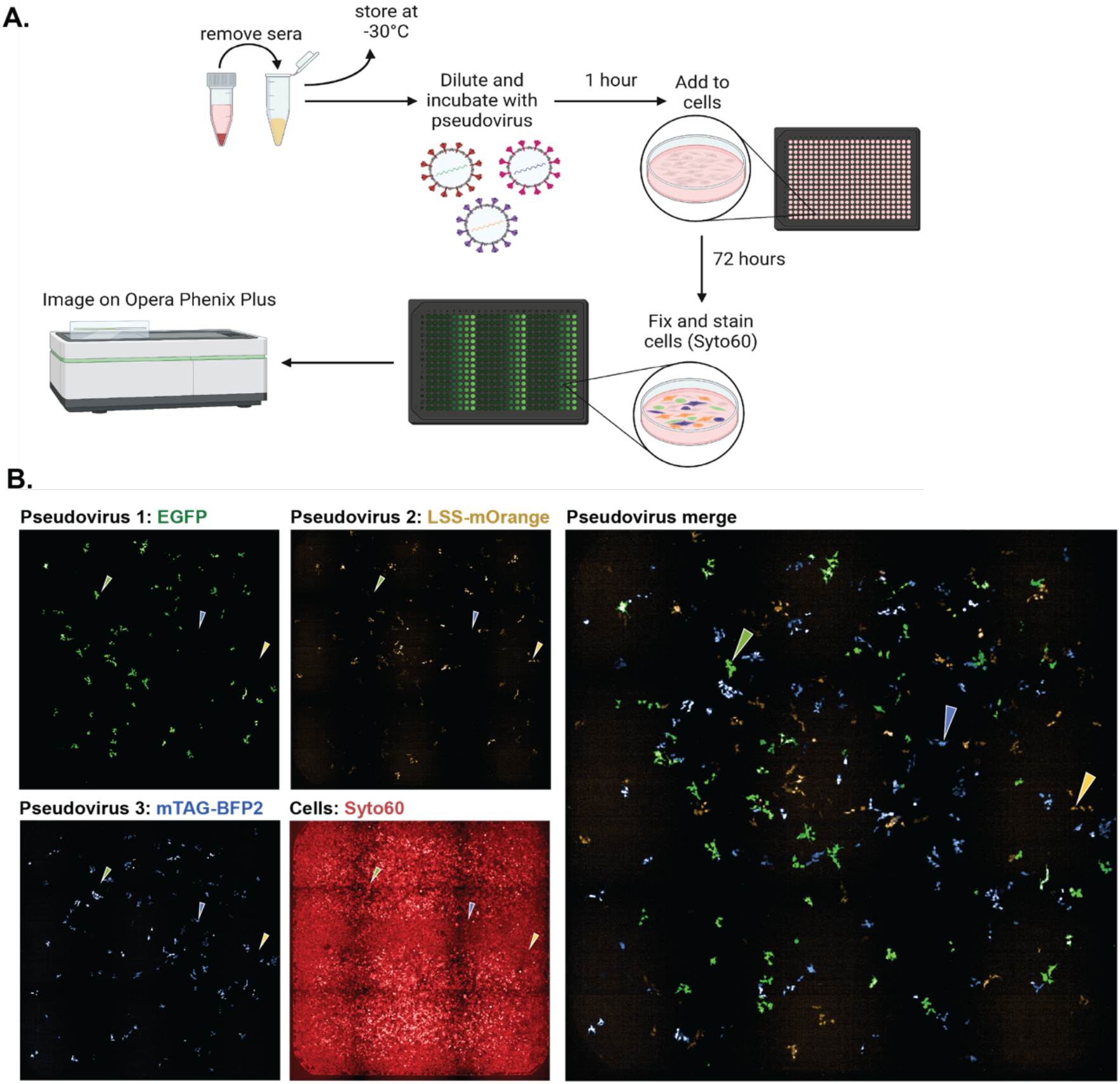
Visualisation of distinct infection of ACE2.HEK293T cells by pseudoviruses expressing different SARS-CoV-2 spike proteins and encoding one of 3 fluorescent reporter proteins. **A**. The workflow of the multiplexed pseudovirus neutralisation assay (pVNT) when used to determine the EC50 titre using small volumes of sera. Four to 6 μL of sera per duplicate (depending on desired dilution factor) is serially diluted before addition of pseudoviruses expressing 3 distinct SARS-CoV-2 spike proteins carrying mRNA for EGFP, mTAG-BFP2 or LSS-mOrange. Sera and pseudoviruses are incubated together at 37°C, 5% CO_2_ for 1 hour before incubation with ACE2.HEK293T cells seeded in a 384-well plate 24 hours prior. Cells are infected by spinoculation 800 xg for 1 hour at 35°C then returned to the incubator for 72 hours. Cells are fixed in 4% paraformaldehyde then washed in PBS and stained with Syto60 for imaging on the Opera Phenix Plus High Content Screening System. Harmony High-Content Imaging and Analysis software is used to determine % infection for each pseudovirus and EC50 titres are calculated by 5PL standard curve interpolation in GraphPad Prism. Image generated in BioRender. **B**. A representative well in a 384-well plate of ACE2.HEK293T cells infected with multiplexed SARS-CoV-2 spike expressing pseudoviruses, coloured arrows indicate cells positive for only one of the three pseudovirus used.

### Optimisation of neutralisation assay parameters to reduce variability

Intra-assay variability has been identified as an issue with both live and pseudovirus neutralisation assays (Kemp et al., 2023; Knezevic et al., 2022; Mukhopadhyay et al., 2022). To identify sources of variation, we examined the contribution of plate location on the final number of cells and percentage transduction in any given well. A 384-well plate was seeded with an equal number of cells per well and then transduced with the same amount of pseudovirus, carrying an EGFP transgene. We examined quantitative and spatial relationships between cell count and the percentage of transduced, EGFP positive cells per well. The mean cell count was 3.7 ×10^4^ (SD = 4.40 ×10^3^) cells per well and the mean percentage of EGFP positive cells was 17.0% (SD = 3.89%) per well. If plate location effects were not present, the proportion of EGFP positive cells was expected to remain constant regardless of the cell count, and values would be randomly distributed across the plate. However, linear regression and a Pearson correlation test revealed a positive correlation between cell count and the percentage of EGFP positive cells per well, which was of moderate strength and statistically significant (r = 0.4959, r^2^ = 0.2459, p < 0.0001) (Figure 2A). Using heatmaps to assess the spatial distribution of cell count (Figure 2A) and percentage transduction (Figure 2B) showed that both measures were generally higher in wells located towards the centre of the plate. Therefore, if the plate layout is asymmetrical (Figure 2D), the nAb titres given by the boxed dilution would be skewed. To minimize the influence of plate effects, a revised, symmetrical plate layout was developed (Figure 2E). The symmetry of this updated layout ensures that wells containing the same antibody concentration experienced consistent conditions. It also features an edge buffer-zone; the outer wells were excluded based on their tendency to contain fewer cells and transduce less effectively (Figures 2B and C). Controls were allocated to the central region of the plate and it was noted that the number of uninfected control wells was equally as important as the number of pseudovirus infected control wells. This is because both are used to set the endpoints of the % neutralisation curves from which the EC50s are interpolated; the uninfected wells establish the level of background noise while pseudovirus only wells represent the maximum level of transduction and its variance. The optimal percentage transduction to aim for with the pseudovirus inoculum was also investigated, using a monoclonal antibody (mAb, J08) derived from a patient infected with SARS-CoV-2 and selected based on the ability to bind the ancestral SARS-CoV-2 spike protein (Andreano et al., 2021). It was found that 5% transduction was optimal to achieve a balance between minimal intra-assay EC50 variation (Figure 2F) and signal values above the background noise (Figure 2G).

**Figure 2.**
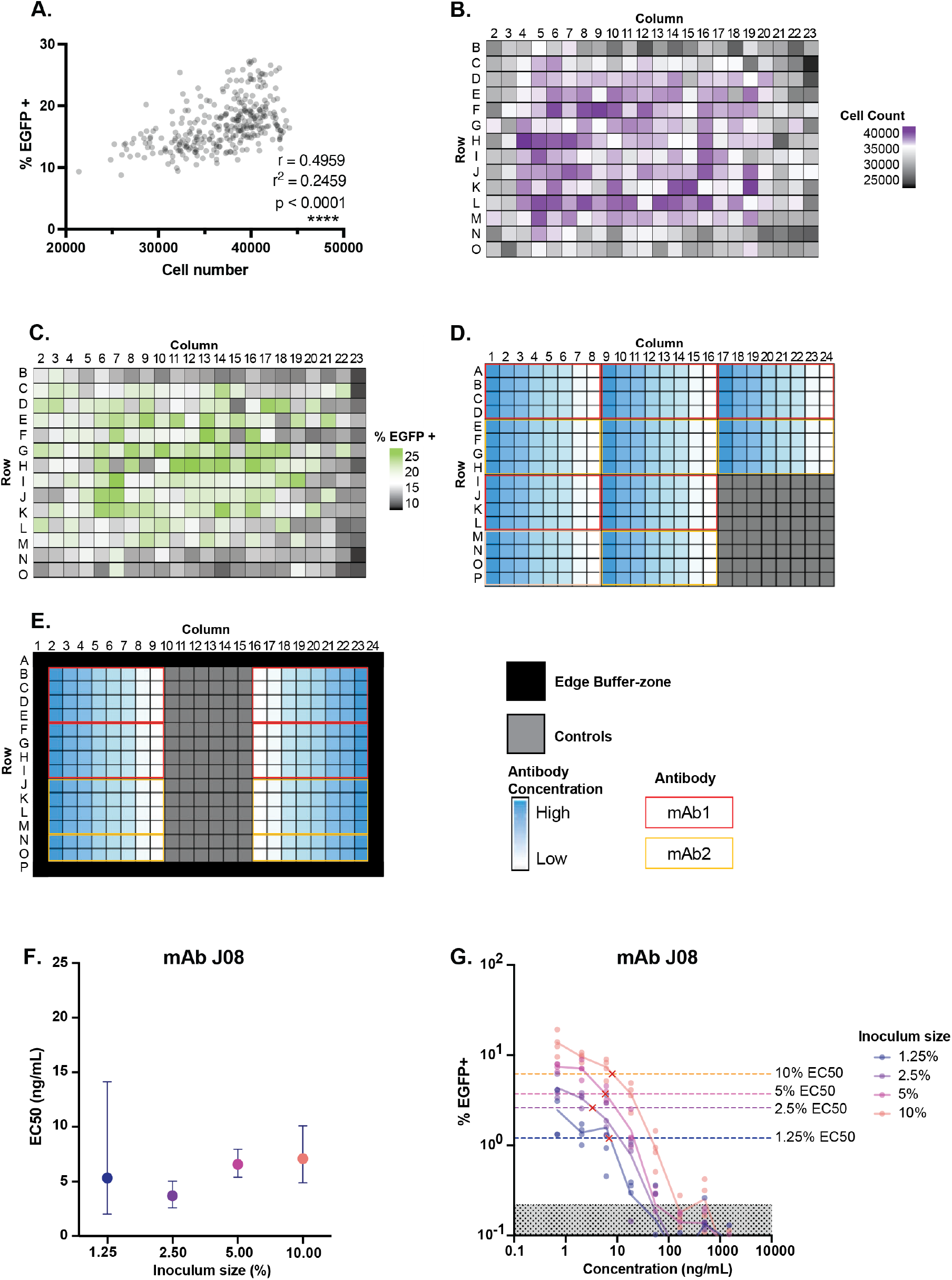
Optimising plate layout and level of pseudovirus transduction to reduce the effect of location and achieve minimal EC50 variation with signal above background noise. An entire 384-well plate of ACE2.HEK293T cells was infected with the same amount of pseudovirus expressing ancestral spike protein carrying an EGFP reporter. **A**. The correlation between the cell count of a well and the percentage of infected cells (%EGFP+) was investigated by linear regression in GraphPad prism. The Pearson correlation coefficient (r), R-squared value (r^2^) and P-value (p) are shown. **B**. Heatmap showing the cell count/well across the plate. Colours are relative to mean cells/well (white). **C**. Heatmap showing %EGFP+/well across the plate. Colours are relative to mean %EGFP+ (white). **D**. The previously used asymmetrical 384-well plate layout, showing arrangement of antibody serial dilutions (yellow and red representing two distinct mAbs) and location of control wells. **E**. Refined symmetrical 384-well plate layout: centrally located control wells, outer wells are excluded, only one antibody tested per plate to minimise inter-plate variation. **F**. Serial dilutions of purified human mAb J08 were prepared and incubated with D614G spike expressing pseudovirus titrated to give 1.25%, 2.5%, 5% or 10% EGFP+ cells. The EC50 (ng/mL) of mAb J08 was determined by pVNT in quadruplicate, EC50 +/- 95% CI are shown. **G**. The curves used to calculate EC50 at each inoculum size (approximate EC50s indicated by red crosses) and where they fall below the background noise (mean of uninfected controls + 2 SD - indicated by shaded area). Coloured dashed lines represent the %EGFP+ corresponding to 50% neutralisation for each inoculum size.

### Neutralisation titres do not differ significantly between single or multiplexed pseudovirus neutralisation assays

Following optimisation of the 384-well plate layout and the pseudovirus inoculum size, the multiplexed pseudovirus assay was used to compare the EC50 titres of mAb J08 in single versus multiplexed pVNTs using pseudoviruses expressing Ancestral, Delta, or BA.1 SARS-CoV-2 spike proteins. mAb J08 showed good potency against all pseudoviruses tested and there were no significant differences in the EC50 calculated for mAb J08 between the single or multiplexed (two or three pseudoviruses) pVNTs (Figure 3A and B, two-way ANOVA, p= 0.0557) To further assess the reproducibility and utility of our single or multiplexed pVNTs, the assays were used to measure neutralisation capacity of patient sera. The patients examined were infected with SARS-CoV-2 early in 2020 in Australia, thus likely were infected with ancestral SARS-CoV-2. Ancestral, Delta or BA.1-specific EC50 neutralisation titres for each patient were determined by single or triple colour pVNT. The percentage neutralisation against dilution factor curves for each patient generated by the single and multiplexed pVNTs were comparable (representative curves in Figure 4A and B). The EC50 calculated were consistent across the single and multiplexed pVNTs (Figure 4C) and were strongly positively correlated across the cohort (r = 0.8598, r^2^ = 0.7393, p < 0.0001) (Figure 4D). Overall, these results demonstrate that pseudoviruses can be multiplexed to accurately measure nAb titres to SARS-CoV-2 variants.

**Figure 3.**
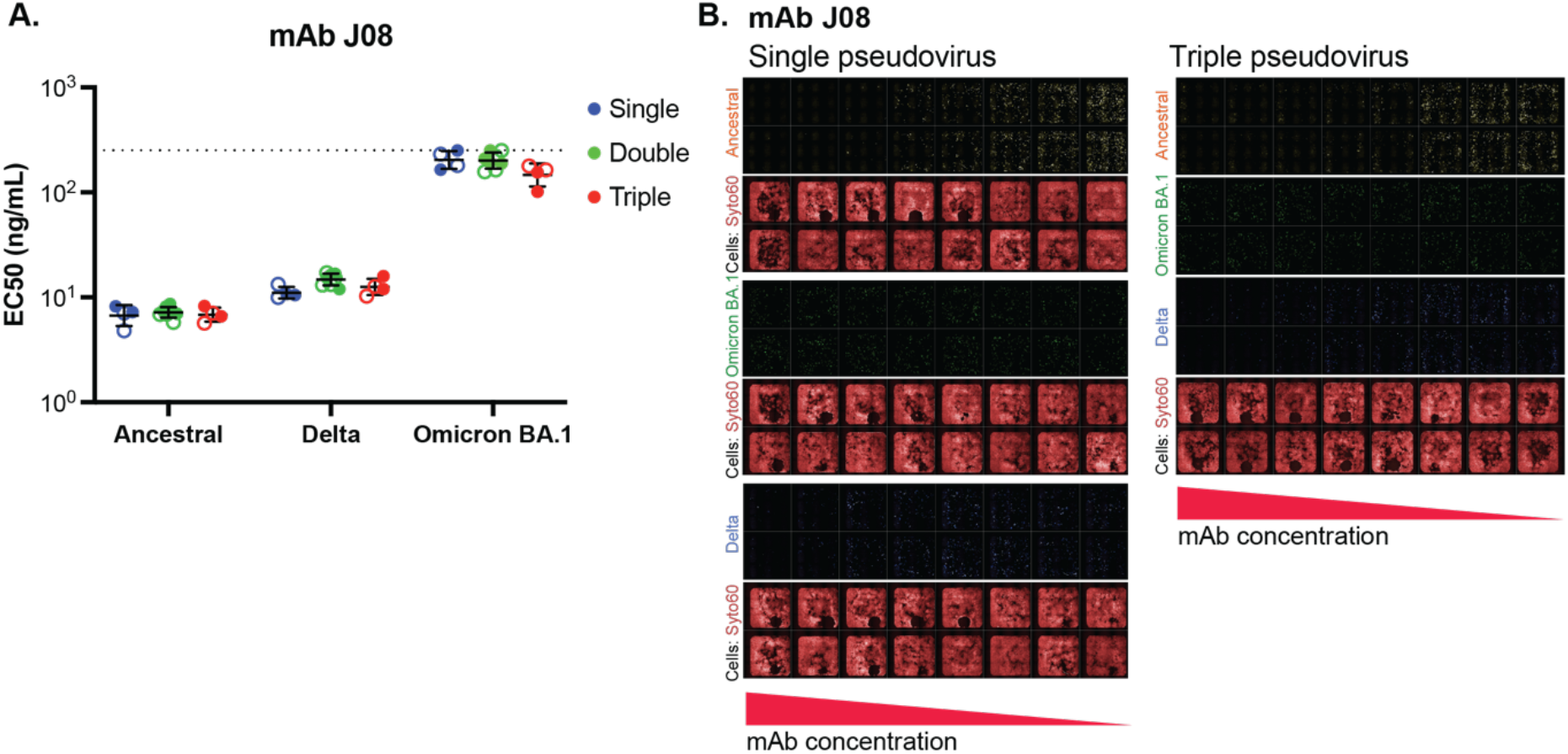
Monoclonal antibody EC50s determined by single pseudovirus assay are not significantly different from those calculated by multiplexed pseudovirus neutralisation assays. **A**. The EC50s of purified monoclonal human antibody J08 against pseudovirus expressing Ancestral, Delta or Omicron BA.1 SARS-CoV-2 spike protein were determined by single, double or triple colour pseudovirus neutralisation assays. Neutralisation by J08 of single and multiplexed pseudoviruses was tested by two users (indicated by open vs. filled circles) in at least two technical replicates each of which were run in duplicate. Geometric mean EC50 +/- geometric SD is shown. Significance was tested for using a two-way mixed ANOVA with Sidak’s multiple comparisons test and none was found. Dashed line indicates the upper assay limit. **B**. Representative images of duplicate wells from single and triple colour assays for mAb J08.

**Figure 4.**
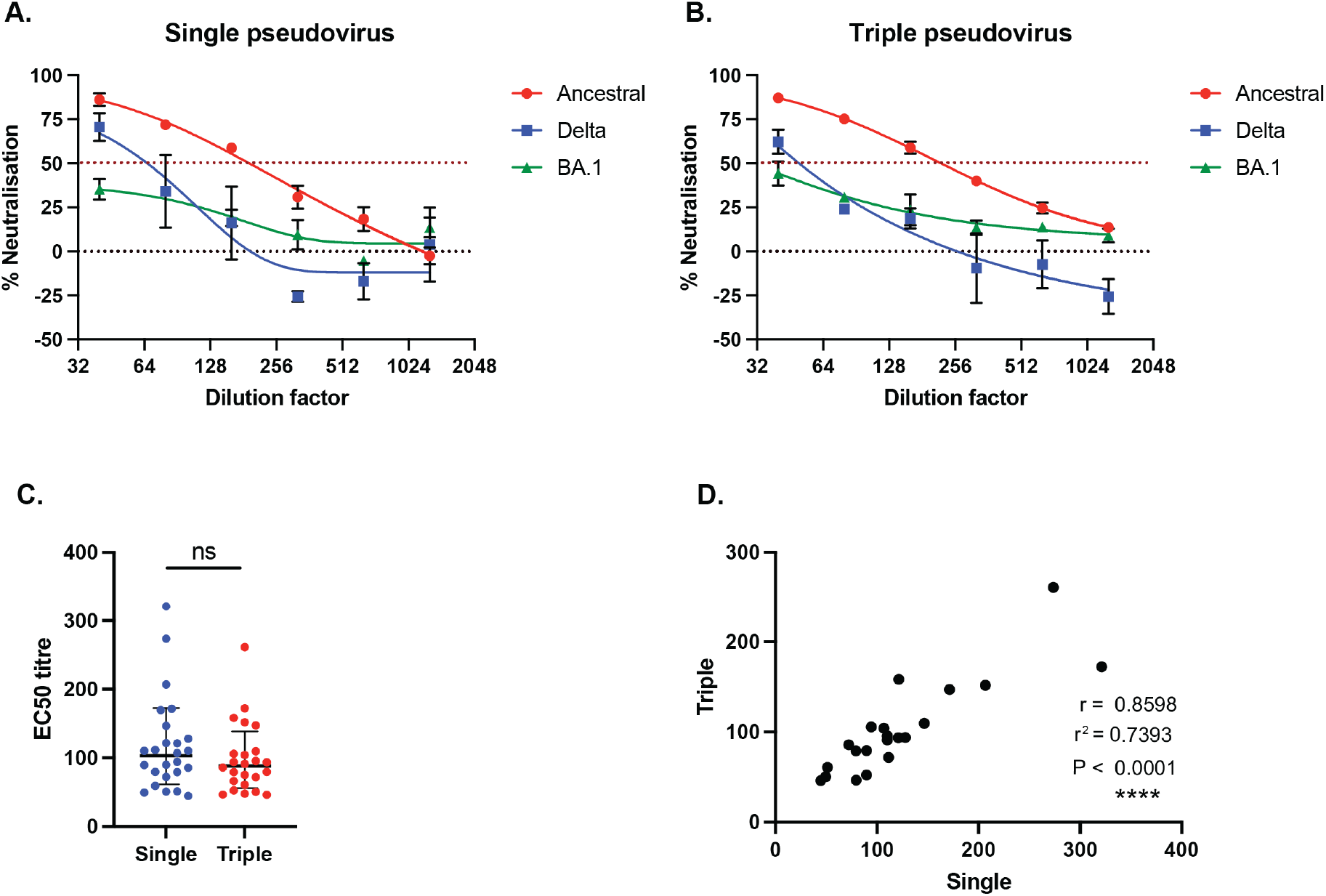
EC50 titres of Ancestral SARS-CoV-2 infected patient sera are comparable between single pseudovirus and multiplexed neutralisation assays. Representative curves used to calculate EC50 for one patient from a cohort of individuals infected in early 2020 **A** single or **B**. triple pseudovirus neutralisation assays (pVNT). Sera was run in duplicate, % neutralisation shown +/-SEM, curves are the result of a 5PL standard curve interpolation in GraphPad Prism 10. **C**. The EC50 neutralisation titres of patient sera calculated from a single pVNT compared to triplex pVNT. EC50s are from a cohort of 10 patients infected in early 2020 whose sera was run against Ancestral, Delta, BA.1, Alpha, Beta and Gamma pseudoviruses singly and across two triplex panels. EC50s falling below the limit of detection not shown. Significance between EC50s calculated by single or multiplexed pVNT was tested for by multiple paired two-tailed t-tests, none was found. **D**. The correlation between the single and triplex pVNT EC50s was investigated by linear regression in GraphPad Prism, Pearson’s correlation coefficient (r), R-squared value (r^2^) and two-tailed p-value are shown.

### Determination of inter-assay variability of the multiplexed pVNT assay

To quantify the inter-assay variability of the multiplexed pVNT, a pool of SARS-CoV-2 nAb positive sera from mice 2 weeks after being vaccinated with a SARS-CoV-2 spike protein in adjuvant using methodology previously described (Counoupas et al., 2022). Sera from these mice was pooled and used to validate the multiplexed pVNT by calculating the coefficients of variation (CV) for EC50s across two, three colour pseudovirus panels in pVNTs performed by two users (Figure 5 A and B). The geometric mean % CV in EC50 across the 8 plates and two independent users who were conducting the pVNTs was 15.9 % (Figure 5C). The CV value is within the 3⋅8–19⋅5 % CV range for SARS-CoV-2 single pVNTs between the seven laboratories of the Coalition for Epidemic Preparedness Innovations (CEPI) Centralized Laboratory Network (Manak et al., 2024). It is also below the 17⋅1–24⋅1% accepted for the gold-standard live virus microneutralisation assay (Manak et al., 2024) and is well within the 8.3%–36.2% CV range for the previously published, multiplexed triple colour human papillomavirus neutralising antibody assay (Nie et al., 2016).

**Figure 5.**
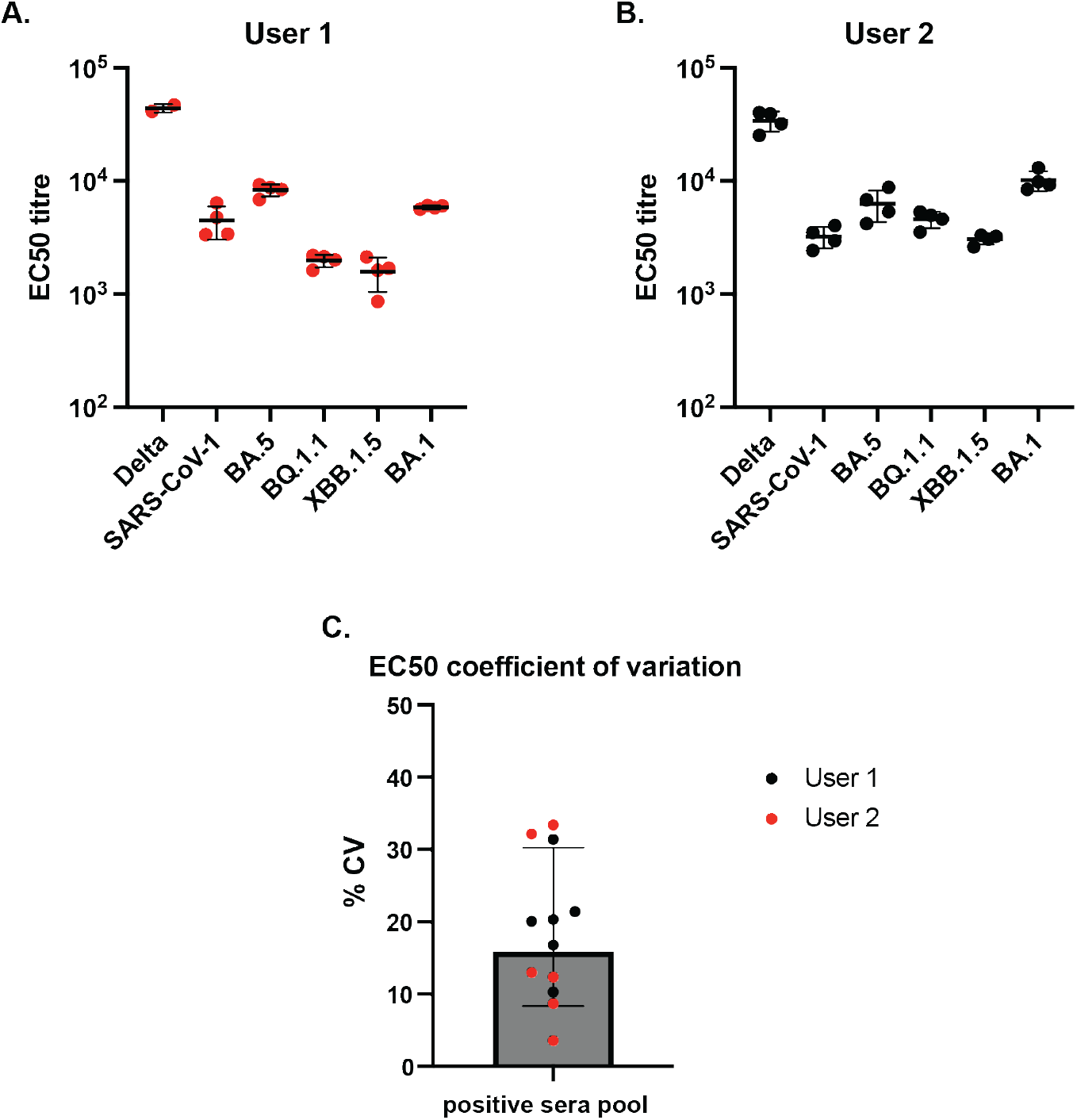
The variation of the optimised, multiplexed pVNT is within the acceptable range. Using two triple colour SARS-CoV-2 spike expressing pseudovirus panels the EC50s of a pool of sera taken from mice vaccinated with chimeric SARS-CoV-2 spike protein were determined. Panel 1: SARS-CoV-1 – EGFP, Delta – mTAG-BFP2, BA.5 – LSS-mOrange. Panel 2: BQ.1.1 – GFP, XBB.1.5 – mTAG-BFP2, BA.1 – LSS-mOrange. Three-fold, 8 point serial dilutions performed in quadruplicate were used to calculate EC50 of this positive, neutralising antibody-containing sera pool which was run on 4 separate plates by 2 users **A**. user 1 and **B**. user 2, geometric mean is shown +/- geometric SD. **C**. The coefficient of variation (% CV) for EC50 for each of the 6 pseudoviruses tested across 8 assay plates was calculated in GraphPad Prism 10. Geometric mean % CV is shown +/- geometric SD.

### Concluding remarks

Looking beyond the 2020 SARS-CoV-2 pandemic, high-throughput systems for the measurement of broadly protective nAbs against a range of coronaviruses will be essential for preclinical screening of vaccine candidates, in the event of a novel spillover event (Cao et al., 2021; Moore et al., 2023; Triccas & Steain, 2024). The multiplexed pVNT described here allows for reliable, simultaneous determination of nAb titres against three distinct coronavirus spike-expressing pseudoviruses in a 384-well plate format. Comparison of monoclonal antibody EC50 titres and variant-dependent fold drops in EC50 titre of human sera between single and multiplexed pVNT assays demonstrated no significant differences, suggesting that the multiplexed pVNT is as robust as the more widely used single pVNT. Thus, the reliability and reproducibility of the assay reported here has utility for widespread use in serological studies, vaccine efficacy assessments, and therapeutic development efforts aimed at combating current and future pandemics.

## Acknowledgments

This work was supported by Coalition for Epidemic Preparedness Innovations (CEPI) as part of the Broadly Protective Beta-Coronavirus Program. We are grateful to the University of Sydney Drug Discovery Initiative and the Sydney Infectious Diseases Institute for support through their seed funding programs. We thank Charles Baily, Centenary Institute, Sydney, Australia for provision of lentivirus packaging and helper plasmids; Stuart Turville, Kirby Institute UNSW, Sydney, Australia for providing SARS-CoV-2 spike plasmid (Ancestral) and ACE-2 293T cells; Nathaniel Landau, NYU Grossman School of Medicine, NY, USA for providing SARS-CoV-2 Delta spike plasmid; and Emanuele Andreano from Fondazione Toscana Life Sciences, Siena, Italy for mAb J08. The authors thank the study participants for their contribution to this research, Angela Ferguson and the RPAVirtual staff who collected the samples. We acknowledge the support of the University of Sydney Advanced Cytometry Facility and the animal facility at the Centenary Institute. Images created with Biorender.com where relevant.

## Notes

### Competing Interest Statement

The authors have declared no competing interest.

